# Neural evidence for decision-making underlying attractive serial dependence

**DOI:** 10.1101/2024.11.18.624176

**Authors:** Jiangang Shan, Jasper E. Hajonides, Nicholas E. Myers

## Abstract

Recall of stimuli is biased by stimulus history, variously manifested as an attractive bias toward or repulsive bias from previous stimuli (i.e., serial dependence). It is unclear when attractive vs repulsive biases arise and if they share neural mechanisms. A recent model of attractive serial dependence proposes a two-stage process in which adaptation causes a repulsive bias during encoding that is later counteracted by an attractive bias at the decision-making stage in a Bayesian-inference-like manner. Neural evidence exists for a repulsive bias at encoding, but evidence for the attractive bias during the response period has been more elusive. We recently [1] showed that while different stimuli in trial history exerted different (attractive or repulsive) serial biases on behavioral reports, during encoding the neural representation of the current item was always repulsively biased. Here we assessed whether this discrepancy between neural and behavioral effects is resolved during subsequent decision-making. Multivariate decoding of magnetoencephalography data during working memory recall showed a neural distinction between attractive and repulsive biases: an attractive neural bias emerged only late in recall. But stimuli that created a repulsive bias on behavior led to a repulsive neural bias early in the recall phase, suggesting that it had already been incorporated earlier. Our results suggest that attractive (but not repulsive) serial dependence arises during decision-making, and that priors that influence post-perceptual decision-making are updated by the previous trial’s target, but not by other stimuli.

## Results

We reanalyzed the data from [1] to investigate the neural correlates of attractive serial dependence. Human participants completed a visual working memory task while their neural activity was recorded with magnetoencephalography (MEG). On each trial (Figure 1A) two gratings were presented sequentially and participants were cued to recall the orientation of one of them. Two different serial biases were found: Participants’ reports were attracted toward the cued item from the previous trial (henceforth “previous target”, Figure 1B; t(19) = 5.8241, *p* < 0.0001). By contrast, when the second sample was cued it was repulsively biased away from the first sample on the same trial (“sample 1”, t(19) = −2.4394, *p* = 0.0247, see [1] for a detailed assessment of behavioral serial biases for each condition). We previously found [1] that the neural biases during the encoding of stimuli failed to fully explain the behavioral biases: in the encoding phase the neural representation of the stimulus was repulsively biased away from sample 1, mirroring the repulsive behavioral bias, but also biased away from the previous target, in contrast to the attractive bias in behavior. Here we re-analyzed the data from this study but focused on a previously unexamined part of the trial: the memory recall period. Our goals were two-fold: First, we wanted to provide neural evidence that the attractive bias is present during the post-perceptual decision-making stage. Second, we wanted to show that this attractive bias depends on context – namely that only the target from the previous trial, but not the other stimulus from the current trial, led to an attractive bias during decision-making.

**Figure 1.**
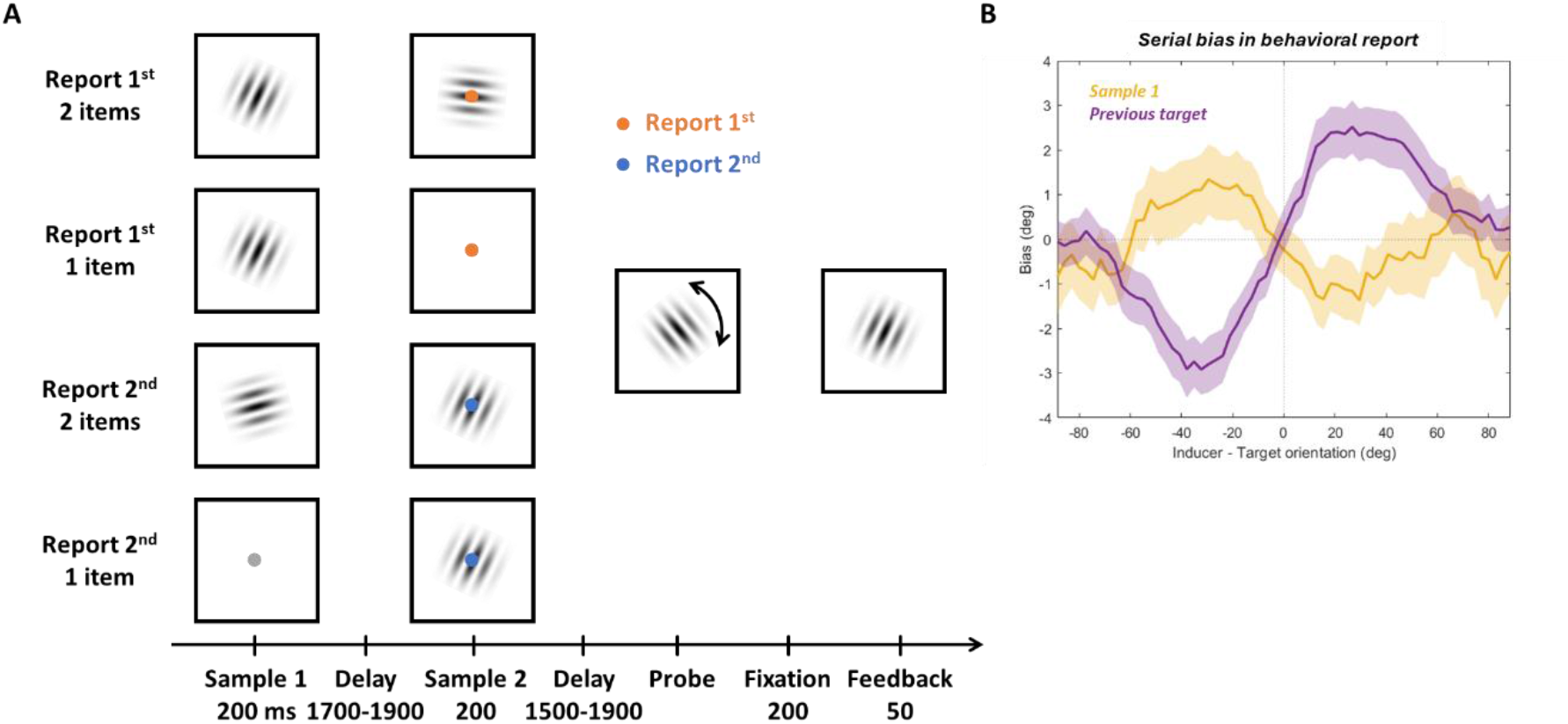
Task design and behavioral results. (A) Two arrays were presented sequentially in each trial. In half the trials two gratings were presented serially (Sample 1 and Sample 2). In the other half of trials, a single grating was presented either at the first (25% of trials) or second (25% of trials) timepoint. A color cue presented at the same time as the second sample indicated which grating the participant should recall. When the probe appeared, participants rotated it to match the orientation of the target item using button presses. The report was registered by pressing a third button. A grating with the correct orientation was presented as feedback. (B) Behavioral serial dependence induced by sample 1 or previous target. The mean signed error of recall errors as a function of the relative orientation of the inducer to the current target. Data was smoothed for visualization purpose only. The smoothing was done by binning the relative orientation of the inducer into 64 evenly spaced, overlapping bins, with each bin containing the 25% of trials closest to its bin center. Shading indicates SEM.

To examine the neural bias in the recall phase we first identified time periods during which the target orientation could be decoded. Multivariate decoding of the target orientation was applied to the MEG data time-locked to the probe onset (see STAR Methods for details). We held out each trial in turn and calculated its neural similarity (using negative Mahalanobis distances) to the average MEG response to all possible stimulus orientations (split into 10 evenly spaced orientation bins, with the held-out trial’s orientation in the center), yielding a tuning curve consisting of the neural similarity to each orientation bin. An unbiased representation of the target should show a peak in the bins closest to 0°. To quantify decoding quality, responses in each bin were transformed to vectors (pointing toward the bin’s orientation and length equal to its similarity to the test trial) and then averaged to a mean vector that was projected onto the 0° vector (Figure 2A). The target orientation could be decoded from 254 to 834 ms after probe onset (Figure 2A left; two-tailed cluster-based permutation test, p=0.00004). We separately time-locked the data to the end of recall (i.e., when participants registered they had finished responding). Target decoding was significant from 163 to 0 ms before the response (Figure 2A right; *p<*0.00001).

**Figure 2.**
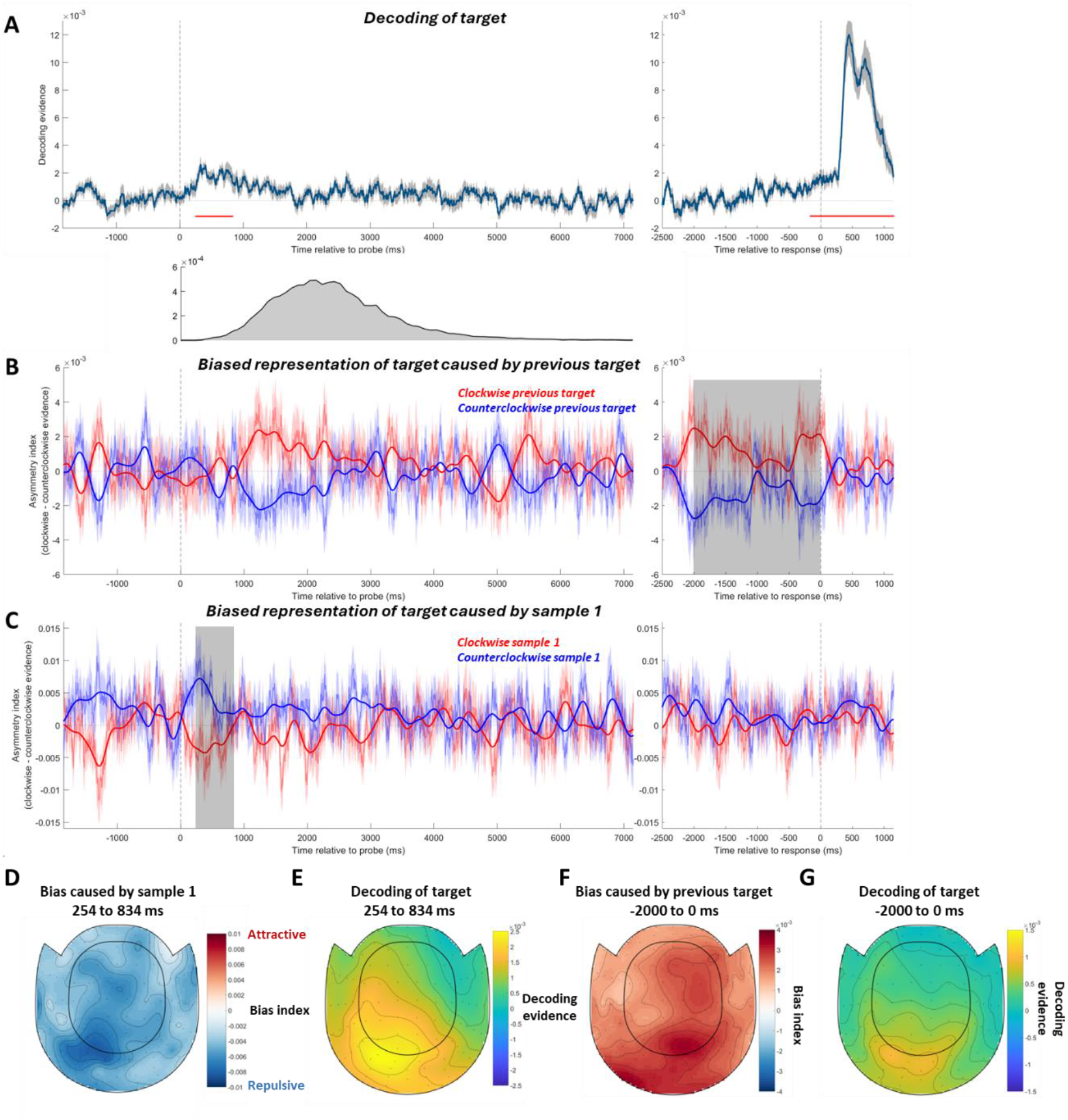
Decoding of the current target and the neural bias on the target induced by sample 1 or the previous target. (A) Orientation decoding of the current target. Decoding evidence of zero would suggest no successful decoding. Red horizontal lines indicate timepoints with significant decoding assessed with cluster-based permutation tests. The plot below shows the distribution of response times across participants and trials. Note that in all probe-locked analyses, the later part of the neural activity contained increasing amounts of noise from irrelevant signals after the current recall period. (B) The bias on the neural representation of current target caused by the previous target. Trials were sorted into two groups according to whether its previous target is clockwise (red line) or counterclockwise (blue line) to the current target. The y-axis shows the mean asymmetry index of them, with a positive number indicating a clockwise-biased neural representation of current target. A smoothed time course is also shown on top (bold lines) for visualization purpose only. The smoothing was done by convolving a Gaussian filter (s.d. = 80 ms) on the raw time course (thin lines). Shadings around the thin lines indicate SEM. The gray-shaded regions indicate time windows with which paired t-tests were conducted. (C) The bias on the neural representation of current target caused by sample 1, with the same plotting conventions as in B. (D) The MEG topography showing which sensors contributed the most to the sample-1-induced repulsive bias, averaged in the time window from 254 to 834 ms after probe onset. The bias index was calculated by subtracting the asymmetry index of counterclockwise inducer trials from clockwise inducer trials. A positive bias index represents an attractive neural bias. (E) The topography showing which sensors contributed the most to the decoding of target in the same time window. (F) The topography of the previous-target-induced attractive bias in the last 2 sec before response. (G) The topography of target decoding in the same time window.

We next tested whether the representation of the current target was biased by previous stimuli by examining whether the neural tuning curve was shifted away from 0°. We took neural similarity to clockwise ([−18°, −72°]) minus counterclockwise orientation bins ([18°, 72°]) as evidence for clockwise vs counterclockwise biases, respectively. The difference between them was calculated as the asymmetry index. A positive asymmetry index indicates a clockwise neural bias.

To investigate the bias induced by the previous target, we compared the asymmetry index of the current target for the trials where the previous target was clockwise vs. counterclockwise to the current target. First, we analyzed the average bias in the time window of significant target decoding (254 to 834 ms after probe onset; Figure 2A). We did not find any bias of the neural representation of the current target caused by the previous target (Figure 2B left; t = 0.4816, *p* = 0.6356). However, a bias seemed to arise later in the recall period (Figure 2B left, starting from ∼1000 ms). As decision-making is a dynamic and variable process (response time distribution in Figure 2A), we investigated the bias locked to the end of the recall period, before participants reached their decisions. In the last 2000 ms before participants completed recall the previous target attractively biased the current target’s neural representation (Figure 2B right; 2000 to 0 ms before response, t(19) = 3.7624, *p* = 0.0013). This attractive bias was still marginally significant when we restricted the analysis to the narrow time window of 163 ms with significant target decoding (163 to 0 ms before response, t(19) = 2.0861, *p* = 0.0507).

To investigate whether sample 1 biased the target during the recall period, we focused on trials where both samples were presented and the 2^nd^ sample was cued to recall. In contrast to the effect of the previous target, sample 1 caused a repulsive bias on the current target’s neural representation immediately upon its reactivation by the probe (254 to 834 ms; t(19) = −3.5613, *p* = 0.0021; Figure 2C). This paralleled its repulsive effect on recall. For the response-locked analysis, in contrast to the previous-target-induced effects reported above, we found no bias induced by sample 1 (last 2000 ms before response: t(19) = −1.0003, *p* = 0.3298; last 163 ms before response: t(19) = 0.3203, *p* = 0.7523).

To further investigate the origins of the attractive and repulsive biases, we conducted a searchlight decoding analysis. The sample-1-induced repulsive bias was most prominent in posterior sensors (Figure 2D). This topography closely resembles the inverse topography of the reactivated target decoding (Figure 2E; Pearson rho = −0.5973, *p* < 0.0001). For the attractive bias found in the response-locked analysis, posterior sensors showed the largest bias (Figure 2F), again similar to the topography of target decoding in the same time window (Figure 2G; rho = 0.668, *p* < 0.0001).

It is possible that the attractive bias we found could be merely driven by the visual response to the probe on the screen. That is, since the participant’s report was attracted toward the previous target and participants rotated the probe stimulus on screen to make this report, it is possible that the attractive bias was actually a sensory effect rather than reflecting a biased memory representation.

To rule out this possibility, we looked directly into the bias of the current target representation driven by the participant’s report on each trial. The logic of this analysis was as follows: On a trial-by-trial basis, the serial bias often diverges from the actual report error because the serial bias only accounts for a relatively small proportion of overall response error variance. We re-sorted the same trials from the analysis of previous-target bias, but this time according to whether the reported orientation (not the previous target) was clockwise or counterclockwise relative to the target orientation. If the stimulus on the screen biased the representation of the target, we should have seen an attractive neural bias towards the direction of the reported orientation. However, no bias caused by the participant’s report survived cluster-based permutation testing. Furthermore, in the time window of the previous-target-induced bias (i.e., 2000 to 0 ms before response), no significant bias was found (Figure 3B; t(19)=0.3208, *p* = 0.7519). During the last 163 ms right before the response (i.e., when the decoding of target was significant), the bias was also non-significant (t(19)=0.1235, *p* = 0.9030).

**Figure 3.**
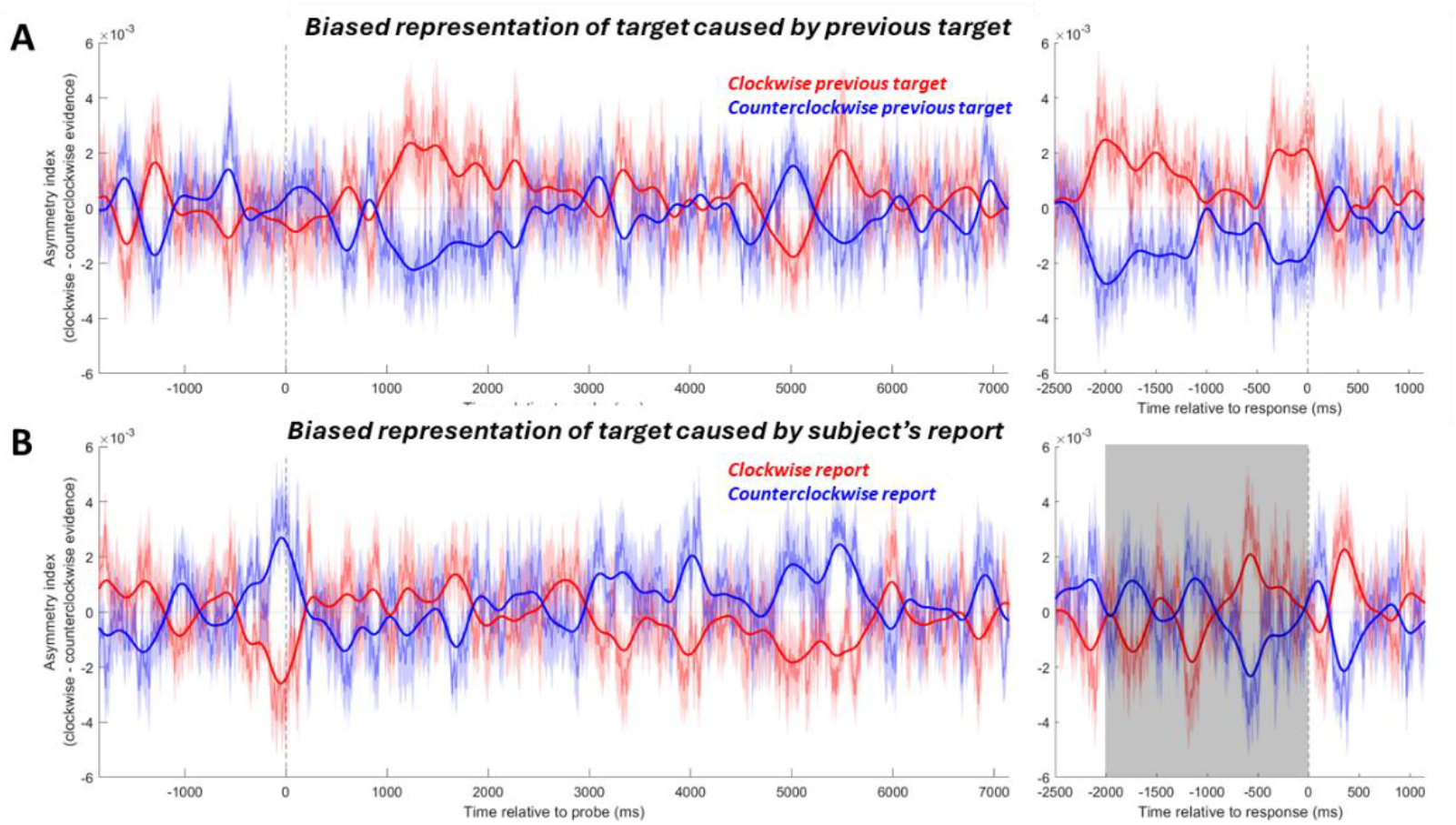
The asymmetry index of representation of current target when trials are sorted according to the relative orientation of the participant’s report made (**B**) and, for reference, the neural bias on the target induced by previous target (**A**).

Furthermore, by comparing the time course and the topography of the previous-target-induced attractive bias and those of the decoding of the participant’s report, we again found evidence inconsistent with this possibility (Figure S1). We conclude that the attractive neural bias towards the previous target was related to biased memory representations and not sensory feedback during recall.

## Discussion

In the present study, we examined how distinct attractive and repulsive serial dependence biases caused by a previous target vs a previous stimulus could arise during working memory recall. Consistent with their respective behavioral effects, we found that sample 1 caused a repulsive bias during the time window when the current target was reactivated by the probe, while the bias caused by the previous target emerged later in the trial and was attractive in nature.

The origin of the attractive serial dependence effect, whether perceptual or post-perceptual, has been a hotly debated question in the field. On the one hand, the continuity field theory of serial dependence proposes that an attractive bias arises during perception and serves to integrate similar stimuli and stabilize noisy perception over time [2]. In support of this view, attractive biases have been observed in early sensory processing [3, 4] and neural activity reflecting the previous target emerges either shortly before [5] or during [6] encoding of a new stimulus. An alternative proposal is that serial dependence of working memory representations develops post-perceptually. For example, a recent model posits that serial bias is the result of a two-stage process: During the encoding of the new stimulus, trial history repulsively biases the encoded representation via sensory adaptation, and during readout of the memory representation, the trial history causes an attractive bias in a Bayesian-inference-like manner [7-10]. In addition to neural evidence for a repulsive bias during the encoding of information [1, 11, 12], an attractive neural bias later in the trial has recently been reported [11, 13]. Our results are consistent with this finding and extended it in two important ways: First, instead of showing the alignment between representational codes for the current- and previous-trial information [11], we examined the neural representation of the target itself and showed direct evidence that this representation was biased toward the previous target in the decision-making stage (as opposed to the memory delay, [13]). More importantly, biases induced by different previous items were examined in the current study. We found results indicating the context-dependent nature of attractive serial bias: only the target from the previous trial, but not the other item from the current trial caused this Bayesian-inference-like attractive bias. While serial dependence is believed to be modulated by attention, such that previously attended information causes a stronger attractive bias [2, 14, 15], our finding suggests different previously attended information may not be incorporated into the prior to the same degree.

Both attractive and repulsive behavioral biases during recall were accompanied by a corresponding neural bias in the recall period, with the sample-1-induced repulsive bias happening quite early and the previous-target-induced attractive bias emerging later. The sample-1-induced repulsive neural bias, at first glance, may seem to be at odds with the two-stage model of serial dependence, since the repulsive bias should happen during the encoding, but not the post-perceptual decision-making stage. However, by closely examining the time courses (Figure 2A and 2C) and the topographies (Figure 2D and 2E) of this repulsive bias and the decoding of the target, it becomes clear that this bias was closely linked to the reactivated representation of the current target. Since the representation of the target was already biased repulsively from sample 1 when it was encoded [1], the repulsive bias found at the very beginning of the recall period could simply reflect the “sensory bias” from the encoding stage instead of emerging for the first time during decision-making.

Why did the previous target and sample 1 exert different biases? From the perspective of Bayesian inference, it suggests that the previous target was incorporated into the prior to inform the decoding of the current target, whereas the sample 1 was not. Behavioral studies on serial dependence have demonstrated that the attractive bias depends on many factors, from the subjective perception of the item [3, 16-18], spatial closeness [15] and shared irrelevant features [19] between the previous and current items, to how participants handled this no-longer-relevant information [20, 21]. In the current study, while participants may have incorporated trial history into the prior by default, for the other memorandum shown in the same trial (i.e., sample 1), they may have had a higher motivation to individuate it and thus excluded it from the prior. Another possibility is that while the participants had actually recalled the previous target, they did not need to recall sample 1. Recall may lead to deeper or prolonged processing and putatively results in a greater impact on the prior. However, we do not believe the attractive bias was driven by an action plan per se, since previous studies have shown that a response to the previous target is not necessary for serial dependence [2, 22, 23], and in the current task the action participants needed to conduct to make the report was independent from the actual orientation of the target (see STAR methods).

Reactivation of the previous information in the current trial has been reported in several previous studies and is believed to be important for attractive serial dependence. Neural representation of the previous stimulus has been found to be un-decodable during the intertrial interval but decodable again around the onset of the current stimulus [24, 25]. Reactivation of previous information has also been observed in the later part of the delay [17] and during the probe period [11, 13]. The strength of the reactivation may correlate with the magnitude of serial bias [5, 17, 26]. In a previous analysis of this dataset, a reactivation of the previous target was found during the encoding of current stimuli [1]. During the probe period we did not see any reactivation of the previous target (Figure S2), in contrast to [11, 13]. A theoretically interesting possibility is that the prior information may influence the decision-making in an activity-silent way, without reactivating itself when participants are making the decision. Further studies at the neural level are needed to study the possible ways in which the prior exerts its influence.

Taken together, by using multivariate decoding of brain activity during working memory recall, we demonstrate an attractively biased neural representation of the current target driven by the previous target. This bias only happened for the target from the previous trial but not the other stimulus previously encoded in the same trial, demonstrating that this post-perceptual decision-making process is context-sensitive. The current study provides evidence for a two-stage model of serial dependence in which adaptation during encoding is overcome by reliance on a temporally smoothed prior during recall of working-memory representations.

## STAR methods

We analyzed the dataset from [1]. 20 participants (11 females) completed a working-memory task where they needed to recall the orientation of one of two gratings presented in the trial (Figure 1A). Each trial started with a black central fixation for 800 ms, followed by the first grating with a random angle presented at the center of the screen for 200 ms. After an interstimulus interval of 1700-1900 ms, the second grating with a random angle was shown for 200 ms. The orientations of the two gratings are independent from each other. Simultaneously with the second grating onset, the central fixation changed color to signal if the first grating or the second grating would be probed. After a delay of 1700-1900 ms, a probe grating was shown on the screen with random starting orientation. Participants were asked to adjust the orientation of the probe to match the orientation of the target grating by pressing two buttons to rotate the probe (Note that this arrangement, along with the random starting orientation of the probe, rendered the action participants needed to conduct to make the report independent from the actual orientation of the target). Responses were submitted by pressing another button. Then, feedback was provided by showing a grating with the correct orientation. Each trial could consist of 1 or 2 gratings, and the probed item could be either the first or the second grating in 2-grating trials. In trials where the first grating was not shown, the fixation dot changed from black to gray to indicate the omission.

Each participant completed 400 trials in total. In 200 trials, both items were presented, with 100 report-first-item trials and 100 report-second-item trials. In 100 trials, only the first item was presented and probed to be recalled. In 100 trials, only the second item was presented and probed to be recalled. Trials were randomly mixed.

### Serial bias on the behavioral level

Serial dependence in the participant’s reports induced by the sample 1 or the previous target was assessed in a model-free way. The averaged signed report error was calculated separately for trials where the inducer is [0°, −45°] or [0°, +45°] relative to the target, and the difference between them is calculated as the index of serial bias for each participant. For the sample-1-induced bias, we focused on trial where both items were presented and the second item was cued to be the target. For the previous-target-induced bias, we used all trials except the first trial in each block, as there was no previous target.

### MEG data preprocessing

To analyzed the MEG activity in the probe period, the data was donwsampled to 250 Hz and re-epoched around the probe onset (from 2 sec before probe onset to 13 sec after probe onset). Epochs were visually inspected to remove trials containing big jumps in activity or non-stereotyped artefacts. Then, an independent component analysis was run on the remaining trials and components related to blinks and cardiac activities were visually identified and removed from the data.

### Multivariate decoding based on Mahalanobis distance

We train the decoder on all non-rejected trials for the decoding of the current target. 367.85 ± 25.609 (mean ± s.d.) trials for each participant were included in this decoding analysis. Response times across all trials ranged from 0.50 to 12.63 sec, and 99% of reports were made 0.70 to 6.20 sec after probe onset (Figure 2A bottom). Thus, we only considered MEG data from 2 sec before probe onset to 7 sec after probe onset in this analysis. For each time point, data from all 306 sensors with a sliding time window of 37 timepoints (i.e., 148 ms) were combined into a vector with 11,322 elements. Principal component analysis was applied on this vector to reduce its dimensionality, maintaining 90% of the variance. We used the Mahalanobis distance to compute the trial-wise distances between the resulting MEG activity patterns, following the practice in [27]. We use a leave-one-trial-out cross validation. Trails are sorted according to the orientation of the to-be-decoded item, for example, the current target: for each testing trial, all training trials were sorted into 10 groups according to the orientation in them relative to the orientation in the testing trial ([−90°, −72°], [−72°, −54°], [−54°, −36°], [−36°, −18°], [−18°, 0°], [0°, 18°], [18°, 36°],[36°, 54°],[54°, 72°], [72°, 90°]). For each timepoint, the Mahalanobis distance between the testing trial and each of the averaged activity patterns from the 10 groups was computed. The 10 resulting distances were mean-centered and the sign was reversed, and then their projections on the 0° vector corresponding to the tested orientation (i.e., each bin’s distance was multiplied by the cosine of the bin orientation) were averaged to yield the decoding evidence. Higher decoding evidence corresponds to stronger and more successful decoding of the item. For the decoding of the previous target, we used this same method but took the target from the previous trial as the to-be-decoded item. The first trial from each block was also excluded from this analysis because there was no previous target. Trials with too short (<0.65 sec) or too long (>11.85 sec) RTs were removed from the response-locked analysis because there is not enough analyzable data at the beginning or the end of the epoch. In the end, 366.95 ± 25.867 trials for each participant were included.

Cluster-based permutation tests with 100,000 iterations were conducted on decoding evidence to assess whether it differs significantly from 0.

### Estimation of neural bias

To test whether the representation of the current target was biased. We looked into the decoding results of the current target and took the mean Mahalanobis distances from channels [−18°, −72°] and [18°, 72°] as clockwise and counterclockwise evidence, respectively. The difference between them was calculated as the asymmetry index. A positive asymmetry index indicates a clockwise neural bias and a negative asymmetry index shows a counterclockwise bias. To investigate the bias induced by the inducer (i.e., previous target or sample 1), trials were separated into two groups according to whether the inducer orientation was clockwise (“clockwise inducer trials”) or counterclockwise (“counterclockwise inducer trials”) relative to the current target. We compared the mean asymmetry index of the current target for these two groups of trials. If the asymmetry index for clockwise inducer trials is higher than the asymmetry index for counterclockwise inducer trials, the neural representation of current target is biased attractively toward the inducer, and vice versa.

For the analyses on biased representation of the target caused by the participant’s report (Figure 3B), we use the same method but take the participant’s report as the inducer. Participants’ reports are known to be subject to systematic biases that are independent of trial history, such as repulsive biases away from the cardinal and oblique axes [16, 28]. These context-independent biases in participants’ reports were corrected first. Following the practice in [16], we fit a sinusoidal function (a sum of three sinusoids with possibly varying frequency, phase and amplitude) on participant’s recall errors as a function of target orientation, and then removed the fit from the recall. Skipping this correction did not change the outcome of the analysis.

For all analyses on neural bias, cluster-based permutation tests with 100,000 iterations were also conducted on the asymmetry index from the two group of trials to assess whether there was a significant difference between them. Due to the higher level of noise in this data, none of the neural bias survived the cluster-based permutation test. Thus, we opted to testing the asymmetry indexes averaged in a time window with paired t-test.

### Searchlight analysis

To test which part of the brain contributed the most to the decoding or the neural bias, we also run searchlight decoding analyses. Iteratively for each one of the 306 sensors, we took this sensor and its 47 most closely adjacent sensors and run the decoding on this smaller set of data. By doing so, we can assess approximately which sensors were more important for the effects of interest. For the topographies of neural bias, the bias index was calculated by subtracting the asymmetry index of counterclockwise inducer trials from the asymmetry index of clockwise inducer trials. Thus, a positive bias index shows an attractive neural bias, and a negative bias index means a repulsive neural bias.

To compare the similarity between topographies, Pearson correlations were calculated.

## Supplementary information

To address the concern that the previous-target-induced attractive bias was actually driven by the rotated probe shown on the screen rather than driven by a higher-order decision-making process, we also conducted multivariate decoding of participants ’reports (that is, to decode the orientation of the report participants actually made instead of the target orientation that they should recall). If the attractive bias was driven by the rotated probe on screen, we should be able to see concurrent decoding of the participant’s report and the attractive neural bias, and the two should have the same posterior-centered topography because they were both stimulus-evoked. Decoding the participant’s report, we found four clusters with significant decoding after the probe onset. For these four time windows, only in one time window a marginally significant attractive bias was found (Figure S1; 1278 to 1814 ms after probe onset: t(19)=1.7479, p = 0.0966). For the other three (Figure S1; 214 to 822 ms, 2522 to 2766 ms after probe onset, and 1055 to 443 ms before participants completed recall), there was not a significant attractive bias (all *p*s > 0.4721).

For the 1278 to 1814 ms time window, searchlight analysis showed the previous-target-induced attractive bias is most prominent in right central sensors (Figure S1C), whereas the concurrent decoding of the participant’s report is most prominent in posterior sensors (Figure S1D). The searchlight topographies were nevertheless positively correlated (similarity between the two topographies measured with Pearson correlation, rho = 0.3978, *p* < 0.0001). While we therefore cannot rule out a contribution of the response to the trend towards a bias in this relatively late phase of the trial, it does not impact our main results, which focused on different time windows that were unaffected by the response.

**Figure S1.**
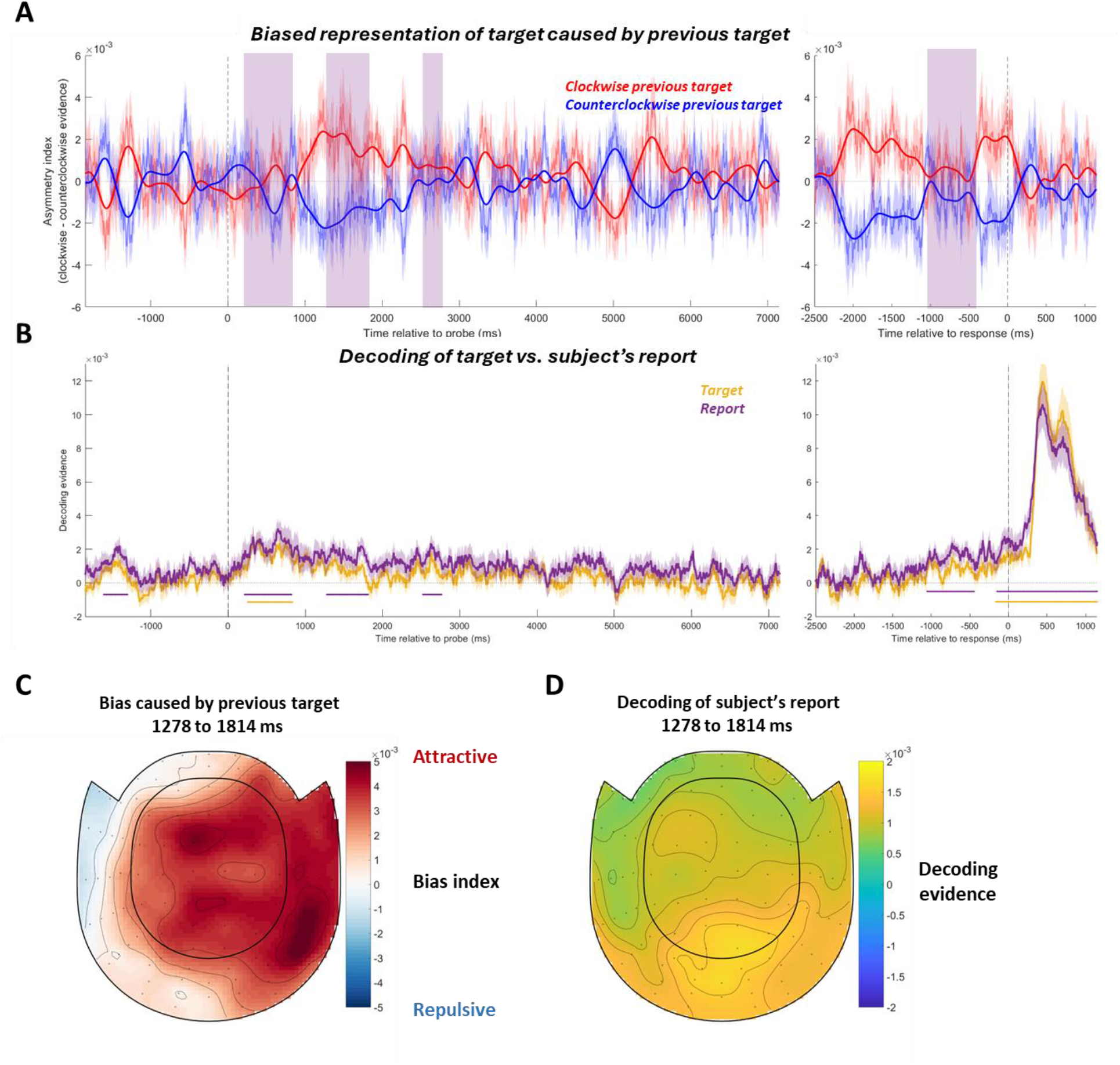
The neural bias on the target induced by previous target (A), the decoding of the current target and the report orientation in the current trial (B), and the topographies of them in the time window where a marginally significant previous-target-induced attractive bias was found (C and D). Yellow and purple horizontal lines in B indicate timepoints where a significant decoding of the target or the report orientation was found, respectively, as tested with cluster-based permutation tests. If the previous-target-induced attractive bias we found was merely driven by the rotated probe on the screen, we should be able to see a significant attractive bias in the time windows where the decoding of the report orientation is significant (purple-shaded regions in A). However, no significant attractive bias was found. In the only time window where a marginally significant previous-target-induced attractive bias was found, the topography of the attractive bias showed important contribution of the right central sensors (C) and the decoding of participant’s report is most prominent in posterior sensors (D). Plotting conventions are the same as in Figure 2.

We interpreted the previous-target-induced neural bias as a genuine shift in the representation of the current target. Alternatively, an apparent attractive neural bias could be the result of the reactivation of the previous target (as has been observed previously, see [5, 24]), in which case we should be able to decode the previous target during the time window where we found the bias. However, in the time window exhibiting the attractive bias no reactivation of the previous target was found (Figure S2, −2000 to 0 ms relative to the report completion, t(19) = −0.1872, *p* = 0.8535). Additionally, no reactivation of the previous target throughout the time course survived the cluster-based permutation test.

**Figure S2.**
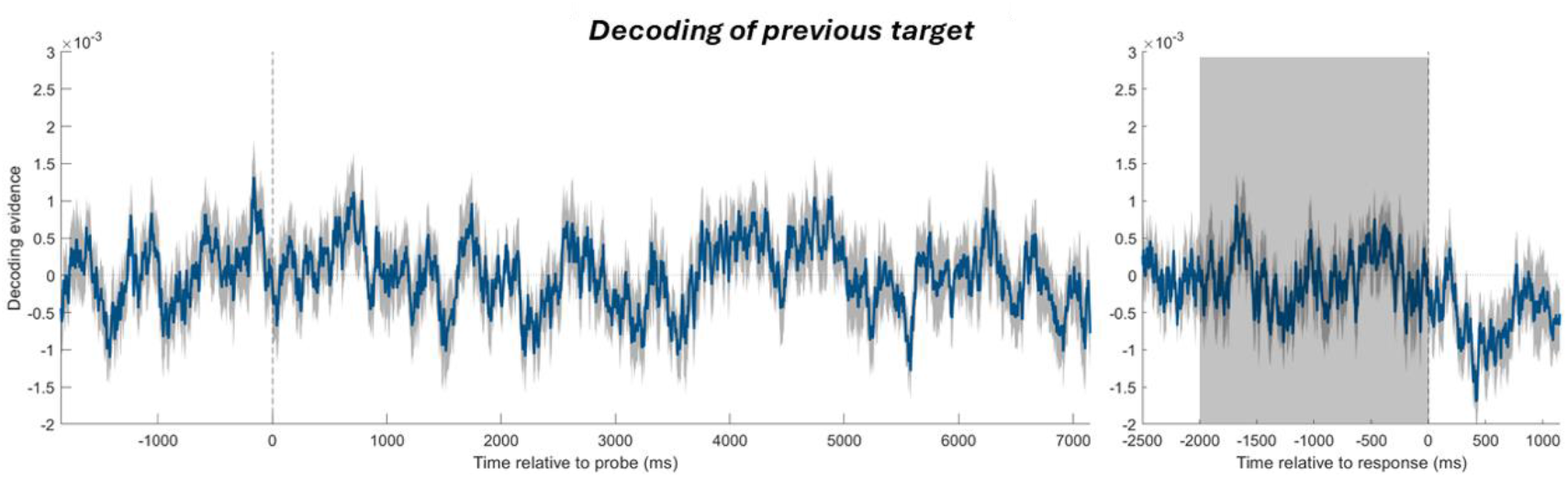
Decoding of the target from the previous trial, with the same plotting conventions as in Figure 2A. Since no significant period of decoding was found with cluster-based permutation test, we also applied one-sample t-tests in the time window where a previous-target-induced attractive bias was found (gray-shaded region). No successful decoding was found in this time window either.

## Acknowledgements

This research was funded by National Institutes of Health (grant number MH131678 to Bradley R. Postle).

